# Plant community associations with environmental variables in a semi-deciduous tropical forest of the Dominican Republic

**DOI:** 10.1101/2020.08.04.235390

**Authors:** José Ramón Martínez Batlle, Yntze van der Hoek

## Abstract

Despite being increasingly threatened by human-induced disturbances, dry forests remain the least studied and protected forest types in the Caribbean region. In contrast to many other forest systems in the world, we have little knowledge of the site-specific variation in vegetation communities within these forests, nor understand how plant species distribution is determined by environmental variables, including among them geological attributes. Here, we assessed the associations between plant communities and habitat types in a semi-deciduous forest of the Dominican Republic. We collected vegetation data from 23 sites within the Ocoa river basin, which we classified into six groups with a Random Forest algorithm, lithology, geomorphology, topography, and last decade history of forest loss as predictor variables. We established three main clusters: one group which encompassed sites with forest over a limestone substrate, four groups of sites with forests over a marlstone substrate with varied degrees of steepness and forest loss history, and one group that gathered all sites with forest over an alluvial substrate. In order to measure the associations of plant communities with groups of sites, we used the indicator value index (IndVal), which indicates whether a plant species is found in one or multiple habitat types, and the phi coefficient of association, which measures species preferences for habitats. We found that 16 species of woody plants are significantly associated with groups of sites by means of their indices. Our findings suggest that the detection of plant species associations with our selection of environmental variables is possible using a combination of indices. We show that there is considerable variation in plant community composition within the semi-deciduous forest studied, and suggest that conservation planning should focus on protection of this variation, while considering the significance and variability of geodiversity as well. In addition, we propose that our indicator groups facilitate vegetation mapping in nearby dry forests, where it is difficult to conduct thorough vegetation or environmental surveys. In short, our analyses hold potential for the development of site-specific management and protection measures for threatened semi-deciduous forests in the Caribbean.

## 1 Introduction

Tropical dry forests are amongst the most threatened tropical ecosystems in the world [26, 40], with 16% of the original area of dry forest remaining in South and Southeast Asia, and 40% in Latin America [40, 45]. In the Caribbean region, dry forests are usually considered relatively resilient to natural disturbances, due to high levels of biodiversity and a high proportion of root biomass, which aids a quick recovery of above-ground parts of plants by rapid absorption of water and nutrients [35]. However, human-induced disturbances decrease this natural resilience and allow for the invasion of alien species [47]. Despite these threats, dry forests remain the least studied and protected forest types in the Caribbean region [5, 47, 49].

Some of the key factors that contribute to the resilience of forest ecosystems threatened by anthropogenic activities, are the structural and compositional diversity of the vegetation, and the local presence of old-growth forest remnants [42, 44]. Such diversity is, in turn, determined by the spatial turnover and distribution of vegetative communities, the heterogeneity of which is linked to spatial patterns among environmental factors [8, 30]. For example, the relationship of climatic variables, topographic metrics and geological traits with the species richness and distribution of vascular plants, highlights the relevance of environmental factors in explaining diversity in a forest ecosystem, which leads to a higher beta-diversity or turnover among vegetation communities [4].

We examined vegetation-environment relationships within a semi-deciduous tropical forest, a type of dry forest, in the Dominican Republic, to assess the association of species with habitats characterized by selected surficial geological attributes as well as and land use, which we group globally as environmental variables. We support recent research aimed to provide evidence of the intimate relationship between living organisms and geodiversity (e.g. abiotic variability), which helps to improve our understanding of the distribution of species [3]. We assessed whether there is spatial variation that can be typified by clusters of plants associated with environmental variables, specifically with lithology, geomorphology, and topography (e.g. slope). Therefore, we hypothesized that some plant species associate with groups of sites characterized accordingly. In addition, we hypothesized that the composition of plant communities is influenced by recent (∼last decade) forest loss, which affected approximately 2% of the study area.

We propose that our findings allow for a better understanding of the spatial variation of vegetation patterns in dry forests in the Dominican Republic, which in turn may inform conservation and management (e.g., restoration) efforts. We detected considerable variation in vegetation composition at relatively small scales sites, for which we identified indicator species [9, 15], even though our study area was small. Since it is logistically challenging to survey all of the dry forests in the Dominican Republic, and conservation planning may focus on protecting as much of the variation as possible, the indicator species we identified will allow us to more easily gain understanding of vegetation patterns in areas with little accessibility. Furthermore, we suggest that our approach can be replicated for other forests and countries in the region, and that our methods can be scaled up to inform regional conservation planning (e.g., maximizing the spatial variation in plant communities included in protected areas).

## 2 Material and methods

### 2.1 Location and data collection

We collected our samples within a *Swietenia–Coccoloba* semi-deciduous forest [22] in the Ocoa river basin of the Dominican Republic (18^°^ 31’ N, 70^°^ 33’ W) (Figure 1). This forest ranges in altitude from almost 300 m to over 800 m, on slopes of varied inclination. The total annual precipitation is 1300 mm, and the mean annual temperature is 24^°^C. The most common lithology is marlstone, but alluvial deposits and limestone are also common.

**Figure 1.**
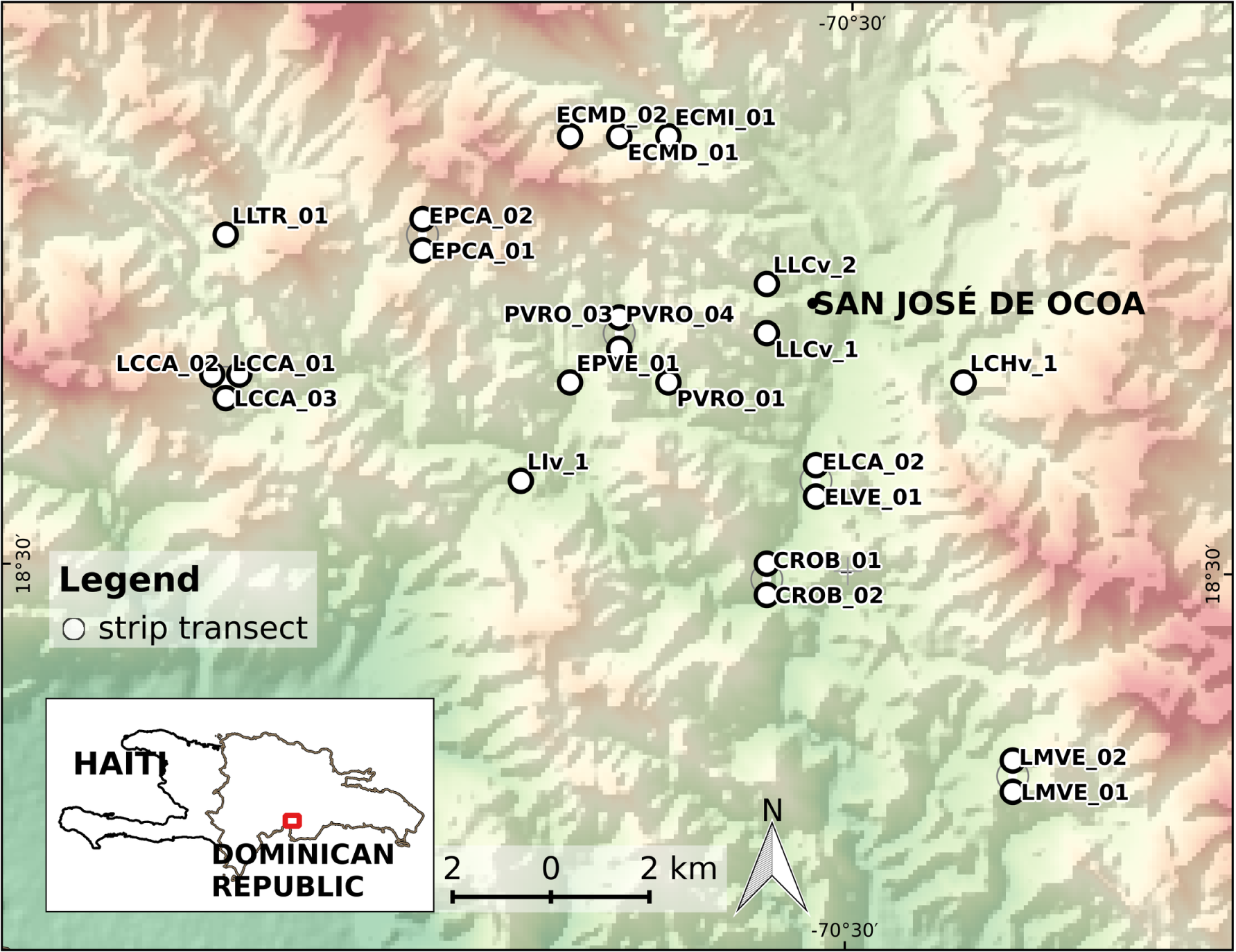
Location map of 23 transects (circles filled white), showing the town of San José de Ocoa. The overlapping points are depicted slightly displaced from their actual positions to ensure that all become visible. The background is a color shaded-relief view (red-white is highland, green is lowland) based on a 30-m SRTM DEM (Ref: NASA LP DAAC, 2000. SRTM 1 Arc-Second Global, https://earthexplorer.usgs.gov/. Published September 2014). The bottom-left inset shows the area in the context of the Dominican Republic.

During two field campaigns (fall 2013 and summer/fall 2014), we collected data using an adapted version of the Gentry transect procotol [20], based on Cámara & Díaz del Olmo [10]. We established 23 randomly placed transects of 50 m ×2 m (100 m^2^), sampling a total area of 2300 m^2^. In our adapted version, we recorded vines, trees and shrubs over 2.5 cm dbh and 1.5 m or more in height. We used height and branching as a criteria for differentiating between trees and shrubs [33]. Thus, we classified as trees the plant individuals at 6 m height and above. Likewise, we classified as shrubs all individuals ramified from the base and below 6 m height.

We digitized vegetation data in `LIBREOFFICE CALC` worksheets [54]. In order to avoid the use of unaccepted names, duplicates and synonyms, we cleaned our data consulting international databases [1, 2] and reviewing previous research [38, 39]. An expert botanist from the National Botanical Garden of Santo Domingo aided us with the identification of the species.

We assessed the completeness of our sample by comparing our numeric species richness with the one obtained in a previous comprehensive study conducted by other research team in a town called Honduras (Peravia province, south of the DR) [46]. This area partly overlaps with the eastern half of the Ocoa river basin, our study area. The authors found 289 plants, including trees, shrubs, lianas and palms. The samples were placed in different types of forest within a range of 300-1400 m above sea level, which included semideciduous forest, but also dry forest, moist forest, cloud forest and others.

Using our abundance data, we estimated the expected numeric species richness with several asymptotic methods available in the `SpadeR R` package [13]. Species richness estimates for our plots ranged from 194 (parametric homogenous model) to 277 species (uncorrected Chao1), which indicates that our observed richness reached between 62% and 89% of the estimated richness. Although we are aware that we sampled across a relatively small study area (few small plots), we were logistically restricted to this sampling design, and deemed our sampling sufficiently complete to justify further analyses.

Using QGIS [50], we placed random points in polygons of rock and landforms in proportions that were representative of the area of the Ocoa Basin covered by each rock/landform type (see Table 1). This allowed us to obtain samples from all rock and landform types found in the region but did not take forest loss into account as a factor of importance for our sampling design. However, during field sampling we obtained local information that there was considerable recent forest loss (after year 2000) in at least two plot locations, something we were able to confirm using a check of maps created by Hansen *et al*. [23]. These authors defined forest loss as a stand-replacement disturbance or a change from a forest to non-forest state, by assessing percent tree cover over time from Landsat imagery. We thereafter decided to utilize this useful, but not a priori considered at the onset of this study, information to conduct preliminary analyses on the impact of forest loss on vegetative community composition. We conducted all fieldwork under a permit issued by the Ministry of Environment and Natural Resources. We requested permission to access private land orally and on-site when required.

**Table 1.**
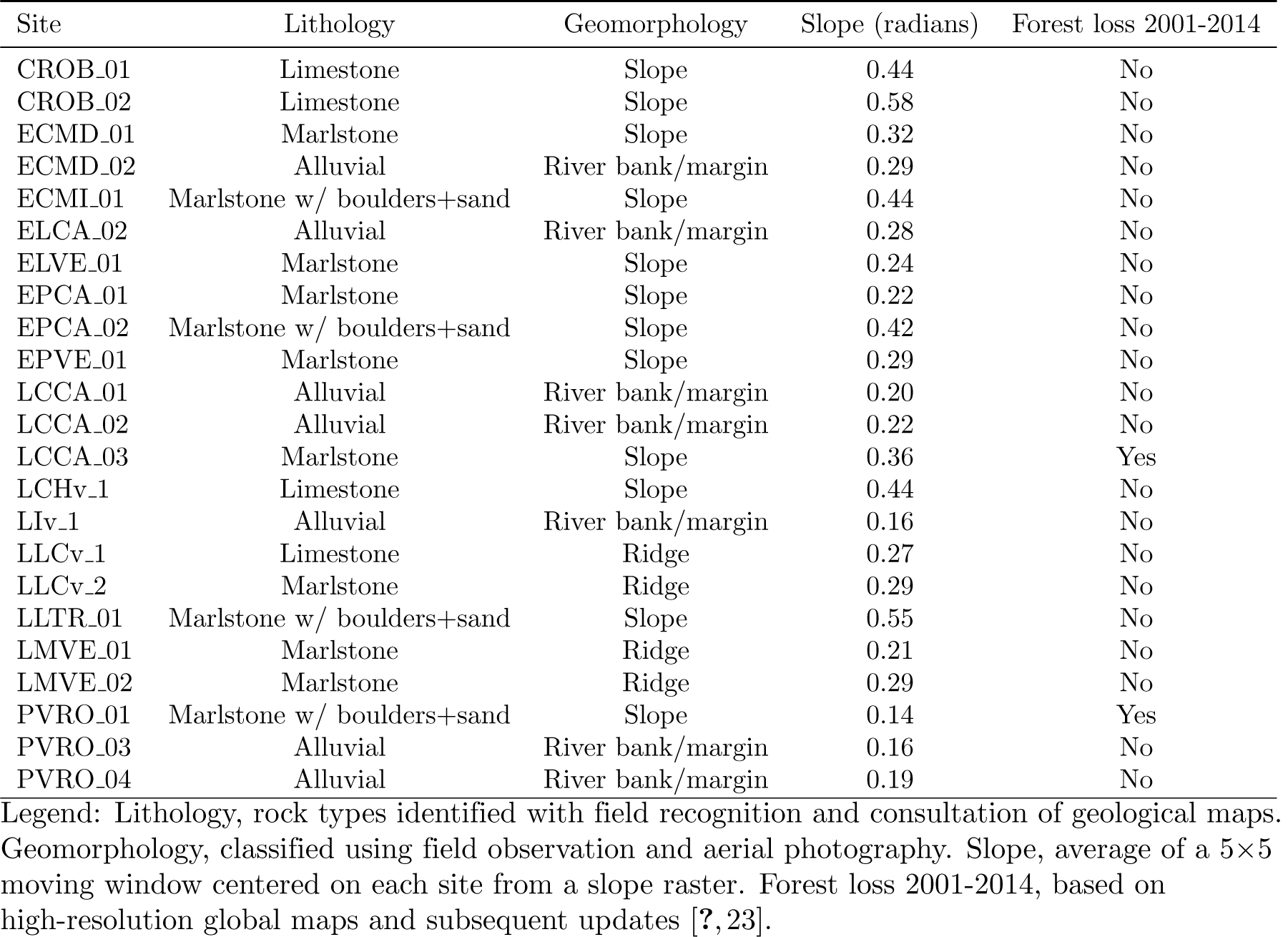
Selected environmental site variables.

Due to the challenging conditions for accessing the forest, we adjusted the predefined location of some of the sites while keeping resemblance with traits of the original points. In addition, to assess the risk of spatial autocorrelation in our data, we conducted a spatial neighbourhood analysis using the actual coordinates of the ultimately selected locations. The average distance between the 23 points was ca. 7.1 km, the minimum was 39 m and the maximum was 19.3 km. It is noteworthy that, from 253 pairwise distances calculated between points, only 9 were below 500 m.

We identified rock types with field recognition and consultation of geological maps (1:50,000 scale) [53], choosing between the following types: limestone (Jura Formation, Middle Eocene), marlstone/mudstone frequently sandy and intercalated with sandstone and boulders (Ocoa Formation, Upper Eocene), marlstone with boulders and sand (Ocoa Formation, Upper Eocene), and alluvial deposits such as boulders, gravels and sand (Quaternary).

We classified landforms qualitatively using field observation and aerial photography [25], choosing between one of three types: slopes, which could be of high or low/medium steepness, river bank/margin, and ridge. As a complement, we also calculated the slope from a terrain processed SRTM 1 Arc-Second Global DEM [41]. To estimate an average angle of slope representative of each entire site, we averaged the slope value of a 5*×*5 pixel moving window centered on each site.

### 2.2 Statistical methods

We conducted all statistical analyses in `R` [51], using the packages `cluster` [36], `factoextra` [28], and `randomForest` [31] to classify sites in groups. For assessing species associations with site groups, we used package `indicspecies` [14]. For zonal statistic we used `raster` [24] and `rgdal` [7], and for data management the packages `reshape2`, `dplyr` and `tidyr` [56–58].

Based on environmental variables, we generated a distance matrix (1-proximity) using the machine learning algorithm *Random Forest* in unsupervised learning mode, a suitable method for variables of mixed types [32]. We configured the algorithm to sample cases with replacement growing 1000 trees, and to use out-of-bag proximity estimation. Subsequently, we used the distance matrix as the source for *AGNES* agglomerative hierarchical clustering algorithm with Ward’s method (agglomerative coefficient of 0.91), then we divided the tree in site groups and characterized them accordingly using environmental traits.

In order to assess the association of species with groups of sites, we calculated the following two indices:

1. *Indicator value index* (hereafter *IndVal*), which assesses the value of a species as a bioindicator. This index was originally proposed by Dufrêne and Legendre [17], as the product of quantity *A*, which measures the *positive predictive value* of a species as an indicator of the site group, and *B*, which measures the *fidelity* or *sensitivity* of the species with the site group. There are variants of the index for presence-absence data and for individual-based data. Also, in the case of unequal group sizes, the authors suggest the equalization of the relative sizes of all site groups. As we collected plant abundance in field campaigns, and our groups are unequally sized, we opted to calculate an “individual-based group-equalized *indicator value index*” using the following formula [9]:

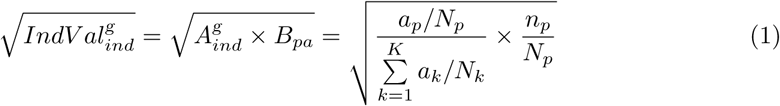

where 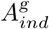 is the mean abundance of the species in the target site group divided by the sum of the mean abundance values over all groups (hereafter *A*_*ind*_), *B*_*pa*_ is the relative frequency of occurrence (presence-absence) of the species inside the target site group, *a*_*p*_ is the sum of the abundance values of the species within the target site group, *n*_*p*_ is the number of occurrences of the species within the target site group, *N*_*p*_ is the number of sites belonging to the target site group, *k* is the number of site groups, *a*_*k*_ is the sum of the abundance values of the species in the *k* th site group, and *N*_*k*_ is the number of sites belonging to the *k* th site group.
2. Pearson’s *phi point-biserial correlation coefficient* (hereafter *r*_*pb*_), in its group-equalized variant, which is suitable for determining the degree of preference of a species for a specific site group among a set of alternative site groups, having individual-based data. We calculated this index according to the following formula (same notation from previous equation) [9]:

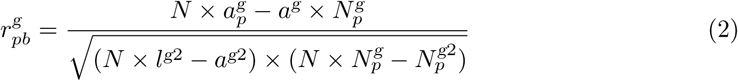

where *l* is the norm of the vector abundances of the species, 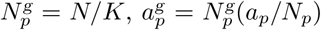, 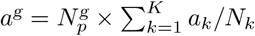 and 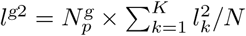. As stated by De Cáceres and Legendre [9], an advantage of *r*_*pb*_ is that it can take negative values, which suggests that species avoid particular environmental conditions. In such a scenario, absences outside the target group contribute to increase the strength of the association, in contrast to *IndVal* which assumes that having fewer or more absences of a particular species outside the target group is not taken into account for measuring the strength of the association.

We used the function `multipatt` from package `indicspecies` to estimate both indices, *IndVal* and *r*_*pb*_. This function computes the value of each index for a site group and also for a combination of them. For each species, the function chooses the site group or the combination of them with the highest association value. Afterward, the best matching patterns are tested for statistical significance of the associations, by means of the permutation test and with *α* set at 0.05.

The permutation test compares an observed statistic with a distribution obtained by randomly reordering the data. Under the null hypothesis of no association, the statistic computed after randomly reassigning the occurrence or abundance values of sites, should be very close to that obtained from unpermuted data. The *p*-value of the test is the proportion of permutations that yielded the same association values than that observed for the unpermuted data. Therefore, we reject the null hypotehsis of no association whether the *p*-value is lower or greater than the significance level [9].

The strategy of randomly reordering the samples varies according to the number of permutations and the random number generator used (the “seed”). Thus, we assessed the sensitivity of our approach by comparing the results of various tests with different combinations of numbers of permutations and seeds. First, we computed 20 tests, by combining 10 different seeds with 10^3^ and 10^4^ permutations each. Afterward, we computed 60 additional tests, by combining 20 different seeds with 10^3^, 10^4^ and 10^5^ permutations each. Overall, we computed 80 *p*-values, and subsequently we summarized them in a conservative approach by keeping the maximum *p*-value from the different permutations tests. Thus, we obtained a short list of species significantly associated with habitats, from which we kept only those species with 4 or more individuals which at the same time were recorded in at least 2 sites.

Finally, as a means of validating our results, we performed a redundancy analysis (RDA), which is a type of canonical ordination aimed to extract the structure of a community matrix in relation to an environmental data set. For this, first we generated a Hellinger transformed matrix from our original community data. Afterwards, we fitted the transformed matrix to the environmental matrix using multiple regression techniques, and summarized the results in a “triplot” showing sites, species and environmental variables in a two-dimensional space. Lastly, we generated a PCA ordination using the Hellinger matrix, and subsequently performed a passive *post hoc* explanation method, by fitting the environmental variables onto the ordination. We assessed the significance of the environmental variables by means of permutation tests. [8, 43].

## 3 Results

During our field campaigns, we recorded a total of 2158 individuals belonging to 172 species, from which 69 are trees, 68 shrubs, 4 palms, 29 vines and 2 cacti. The vast majority are native (n=130, 76%) and endemic species (n=34, 20%). The eight most abundant species represented almost one-third of the abundance, which included *Coccoloba diversifolia* and *Randia aculeata*. We collected an average of 93 individuals per site, with a maximum of 175 individuals (LCCA 03) and a minimum of 47 individuals (LLTR 01). The average numeric richness per site was 28 species, with the richest site reaching 44 species (LCCA 02) and the poorest just 16 species (CROB 02).

### 3.1 Site groups

We classified our 23 sites in six groups, which we characterize below according to environmental traits (see Figure 2):

**Figure 2.**
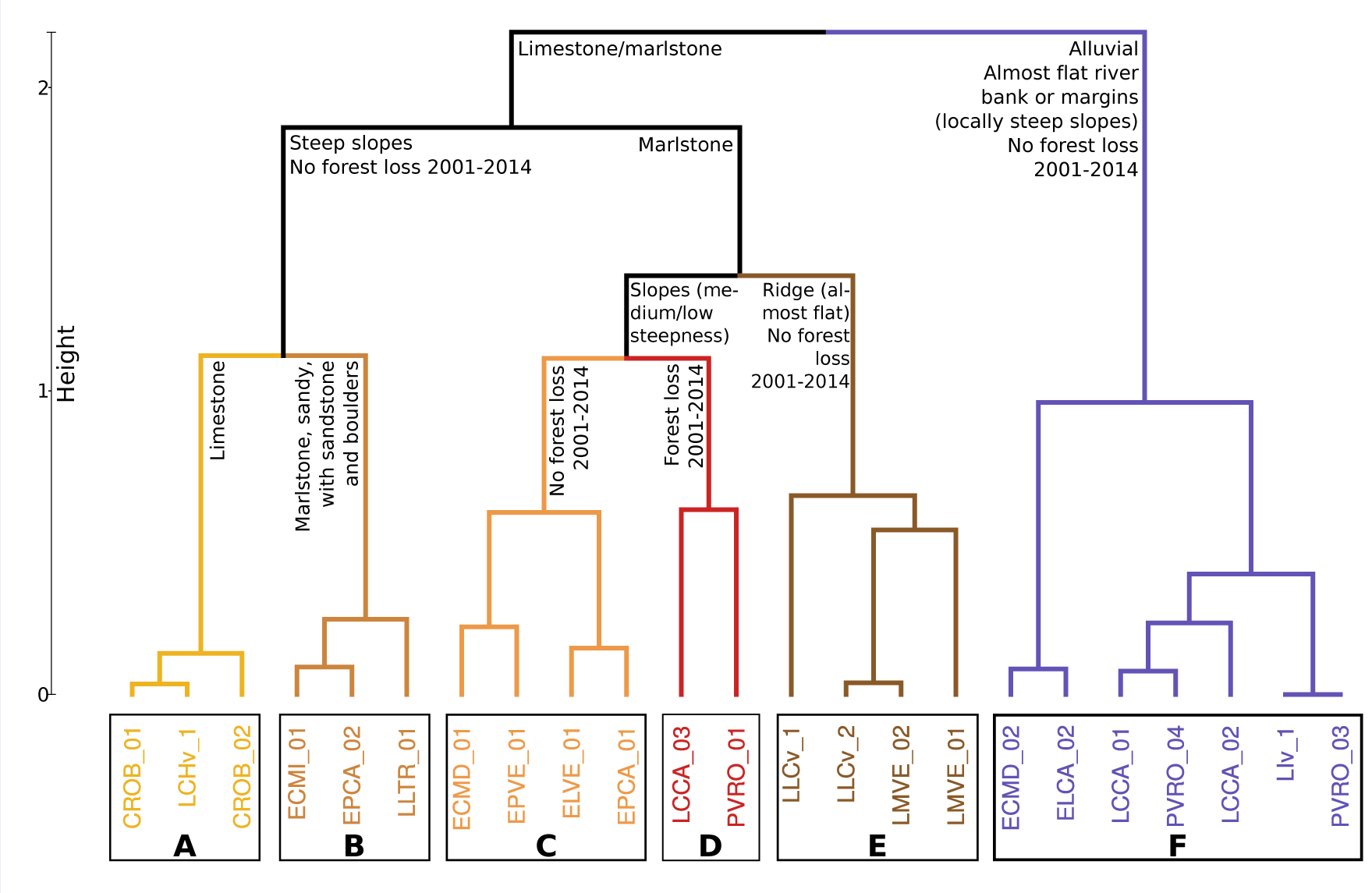
Dendrogram of groups of sites based on selected environmental variables. Each group is indicated with bold letters from A to F. See text for details.

1. Group A. Forests on steep slopes consisting of limestone, no forest loss reported from 2001 through 2014.
2. Group B. Forests on steep slopes consisting of sandy marlstone, intercalated with sandstone and boulders, no forest loss reported from 2001 through 2014.
3. Group C. Forests on medium to low steepness slopes consisting of marlstone, no forest loss reported from 2001 through 2014.
4. Group D. Forests on medium to low steepness slopes consisting of marlstone, occasionally with boulders and sands, forest loss reported from 2001 through 2014.
5. Group E. Forests on almost flat ridges consisting of marlstone, no forest loss reported from 2001 through 2014.
6. Group F. Forests on almost flat river banks or margins (locally steep slopes) consisting of alluvial deposits, no forest loss reported from 2001 through 2014.

### 3.2 Indicator value index, *IndVal*

*IndVal* calculations suggest that 11 species are significantly associated with groups of sites (Table 2). Five species showed significant association with only one group of sites, and the the other six species showed a strong association with combinations of groups of sites.

**Table 2.**
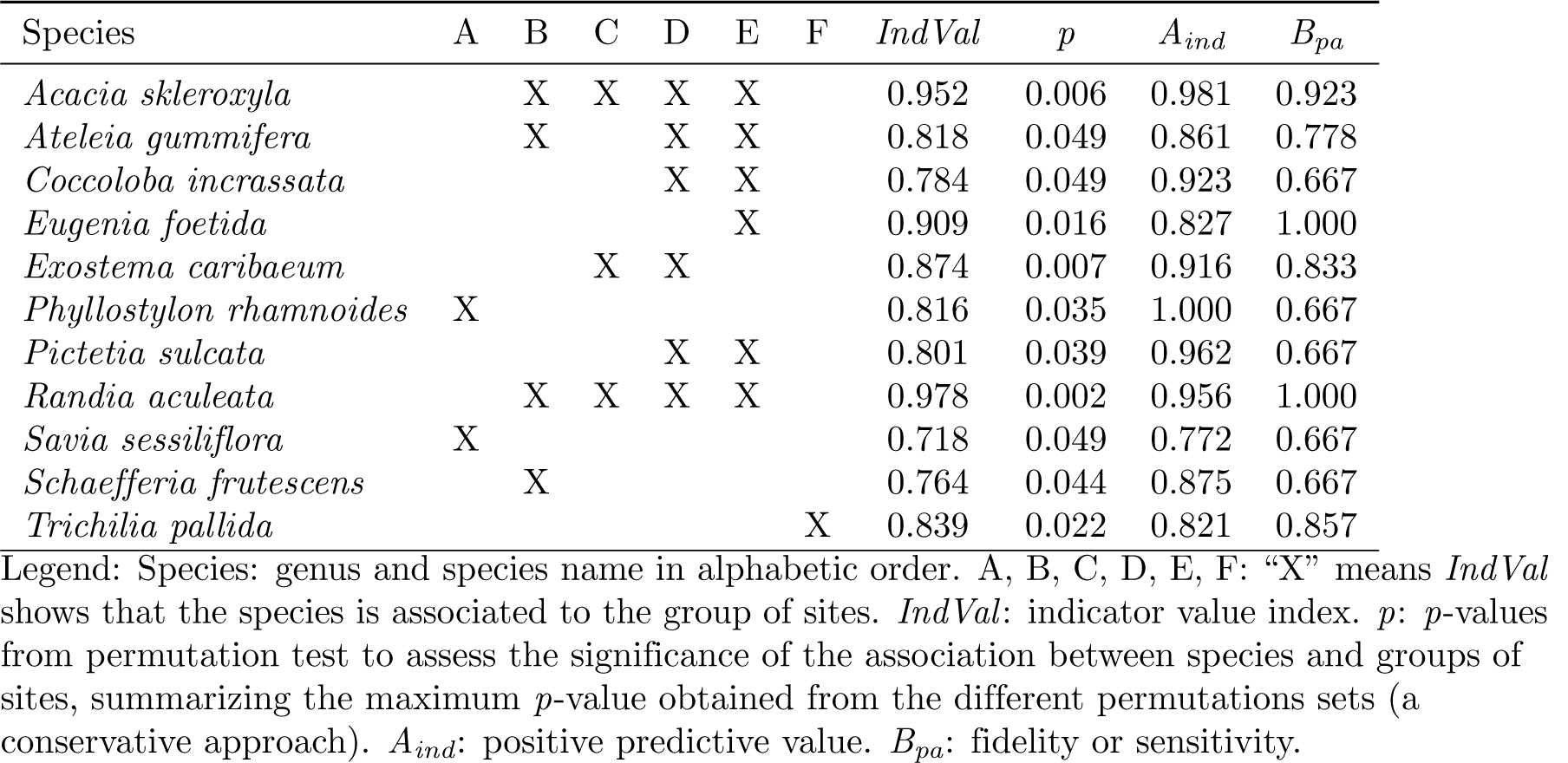
*IndVal* calculations for 11 significantly associated species with groups of sites.

We highlight that *Phyllostylon rhamnoides* and *Savia sessiliflora* are suitable indicators for group A (forests on steep slopes consisting of limestone). Two species, *Acacia skleroxyla* and *Randia aculeata*, showed a strong association with forest growing on marlstones in different geomorphological positions. Also, we found that *Eugenia foetida* is a good indicator of forests on flat marlstone ridges (group E), and tests showed that *Schaefferia frutescens* is significantly associated with group B, which comprises forests on steep slopes consisting of sandy marlstone, intercalated with sandstone and boulders. Moreover, we found that *Trichilia pallida* is a suitable indicator of group F, which are forests on almost flat river banks or margins (locally steep slopes) consisting of alluvial deposits.

### 3.3 Pearson’s phi point-biserial correlation coefficient, *r*_*pb*_

Ecological preference, measured by *r*_*pb*_, showed that 13 species significantly associate with one group of sites or a combination of them (see Table 3). From this, seven species are shared between this list and that of the *IndVal* calculations, also showing preference for the same groups of sites with which they were associated through *Indval*. In addition, we highlight *Ateleia gummifera, Coccothrinax argentea* and *Leucaena leucocephala*, associated with group D, which represents forests on marlstone with tree cover loss reported between 2001 and 2014. Lastly, we highlight two other species, *Hura crepitans* and *Picramnia pentandra*, which showed ecological preference with group F.

**Table 3.** *r*_*pb*_ calculations for 13 significantly associated species with groups of sites.

| Species | A | B | C | D | E | F | *r<sub>pb</sub>* | *p* |
| --- | --- | --- | --- | --- | --- | --- | --- | --- |
| *Ateleia gummifera* |  |  |  | X |  |  | 0.752 | 0.022 |
| *Coccoloba incrassata* |  |  |  |  | X |  | 0.696 | 0.048 |
| *Coccothrinax argentea* |  |  |  | X |  |  | 0.990 | 0.007 |
| *Eugenia foetida* |  |  |  |  | X |  | 0.796 | 0.009 |
| *Exostema caribaeum* |  |  | X | X |  |  | 0.727 | 0.032 |
| *Hura crepitans* |  |  |  |  |  | X | 0.694 | 0.046 |
| *Leucaena leucocephala* |  |  |  | X |  |  | 0.683 | 0.046 |
| *Phyllostylon rhamnoides* | X |  |  |  |  |  | 0.706 | 0.035 |
| *Picramnia pentandra* |  |  |  |  |  | X | 0.668 | 0.049 |
| *Pictetia sulcata* |  |  |  | X | X |  | 0.656 | 0.049 |
| *Samyda dodecandra* |  |  |  | X |  |  | 0.708 | 0.028 |
| *Schaefferia frutescens* |  | X |  |  |  |  | 0.703 | 0.045 |
| *Trichilia pallida* |  |  |  |  |  | X | 0.700 | 0.039 |

Legend: Species: genus and species name in alphabetic order. A, B, C, D, E, F: “X” means *r<sub>pb</sub>* shows that the species is associated to the group of sites. *r<sub>pb</sub>* stands for *phi* point-biserial correlation coefficient.

### 3.4 Summary of species associated with site groups

Overall, 16 species are significantly associated with groups of sites by means of *IndVal* and/or *r*_*pb*_ indices (see Table 4). We separated three sets of species based on whether the association to site groups was detected by both indices, or only by one of them:

**Table 4.** List of species significantly associated with one or more groups of sites by means of *IndVal* and/or *r*_*pb*_.

| Species | *IndVal* | $r_{pb}$ |
| --- | --- | --- |
| *Acacia skleroxyla* | X |  |
| ***Ateleia gummifera*** | X | X |
| ***Coccoloba incrassata*** | X | X |
| *Coccothrinax argentea* |  | X |
| ***Eugenia foetida*** | X | X |
| ***Exostema caribaeum*** | X | X |
| *Hura crepitans* |  | X |
| *Leucaena leucocephala* |  | X |
| ***Phyllostylon rhamnoides*** | X | X |
| *Picramnia pentandra* |  | X |
| ***Pictetia sulcata*** | X | X |
| *Randia aculeata* | X |  |
| *Samyda dodecandra* |  | X |
| *Savia sessiliflora* | X |  |
| ***Schaefferia frutescens*** | X | X |
| ***Trichilia pallida*** | X | X |

Legend: *IndVal* and $r_{pb}$ : “X” indicates the species is significantly associated with one or more groups of sites, whether by means of only one of the indices or by both of them simultaneously. Species **highlighted in boldface** are associated to site groups by both indices, which suggests a strong association.

1. Eight species that are significantly associated with site groups by both indices: *Ateleia gummifera, Coccoloba incrassata, Eugenia foetida, Exostema caribaeum, Phyllostylon rhamnoides, Pictetia sulcata, Schaefferia frutescens* and *Trichilia pallida*. Thus, these species are indicators and also have significant preference for the habitat types represented in site groups, meaning that associations are likely to be strong.
2. Three species that are associated with site groups by means of *IndVal* only: *Acacia skleroxyla, Randia aculeata* and *Savia sessiliflora*. These species are indicators of one or more of their associated site groups.
3. Five species that are associated with site groups by means of *r*_*pb*_ only: *Coccothrinax argentea, Hura crepitans, Leucaena leucocephala, Picramnia pentandra* and *Samyda dodecandra*. These species show an ecological preference to one or more of the habitats represented by their associated site groups.

### 3.5 Canonical ordination analyses

We performed a redundancy analysis (RDA), a type of constrained community ordination that incorporates the explanatory variables directly in the ordination process. This technique is suitable for extracting the structure of the composition data set (e.g. the community matrix) that are related to the environmental variables. We found that one third of the total variance explained by the RDA corresponds to the constrained fraction, which is a high value considering the high complexity of our community matrix.

We represented the RDA using a triplot, a graph that features three types of entities in a single ordination space: sites, response variables (e.g. species) and explanatory variables, which in this case are environmental variables (see Figure 3). The triplot resembles the six clusters identified by the AGNES agglomerative hierarchical clustering algorithm. In addition, the species correlated with the sites are the same highlighted as associated by both *IndVal* and/or *r*_*pb*_ indices to any of the groups previously identified. This is an expected result, since the RDA incorporates the environmental variables in the ordination process.

**Figure 3.**
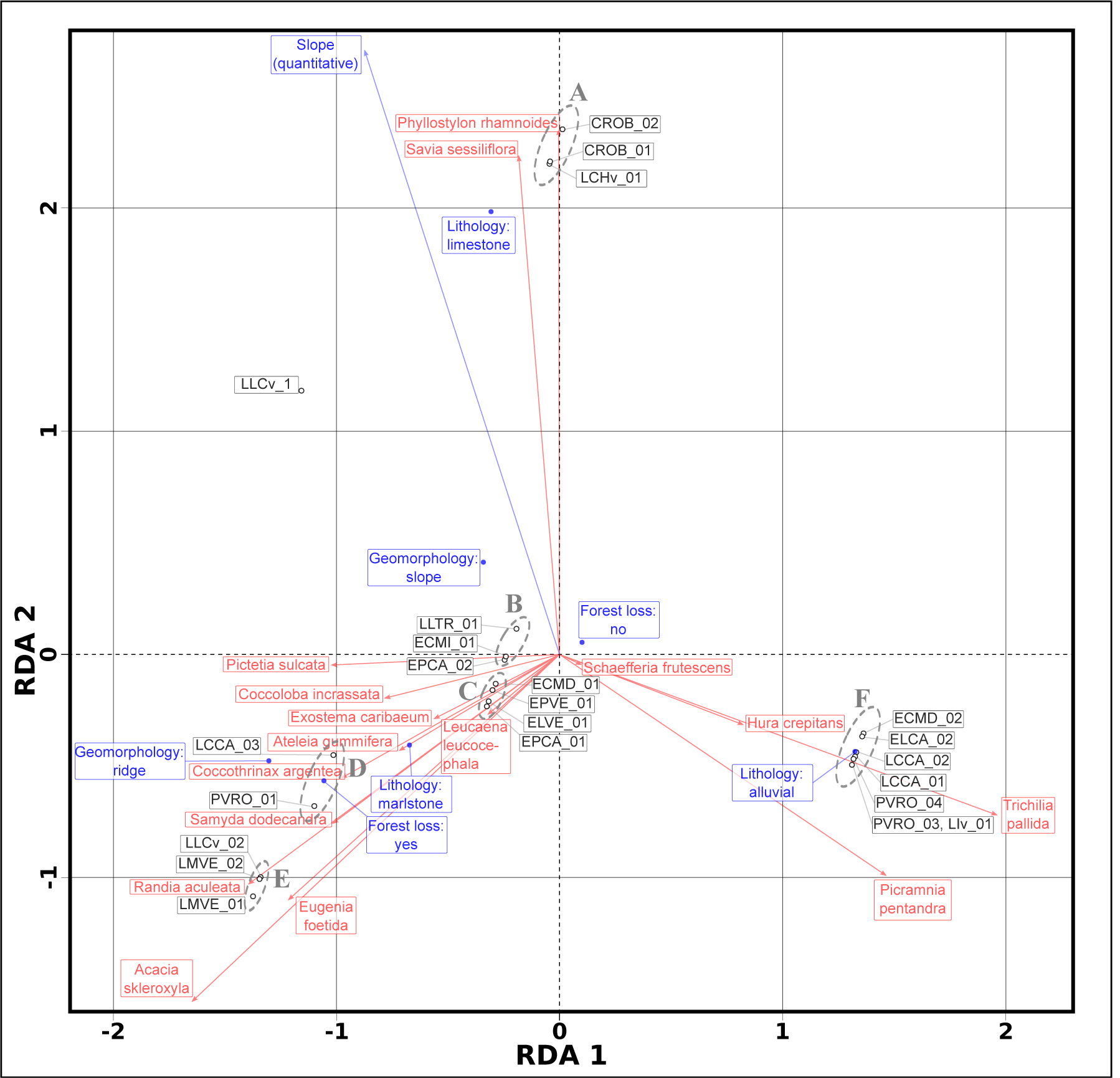
Triplot of the Hellinger-transformed abundance community matrix, constrained by the variables (blue outlined rectangles) lithology, geomorphology, forest loss 2001-2014 and slope. Though we used the entire community matrix for computing the RDA, we only plotted the 16 species associated by means of *IndVal* and *r*_*pb*_. This is a scaling 2 triplot, where angles between species and explanatory variables, and angles between species themselves and variables themselves, are interpreted as correlations. The labelled (A to F) ellipses with grey dotted border enclose site groups defined by the AGNES agglomerative hierarchical clustering algorithm. See text for details.

We also tested the significance of the environmental variables in relation to a PCA ordination of the Hellinger transformed community matrix. The results of the fitting function and the permutation tests suggest that both lithology and geomorphology are significant factors in our sample. However, forest loss history and terrain slope resulted non-significant variables according to the tests (Table 5).

**Table 5.** Regression results from fitting the environmental variables onto a PCA ordination of a Hellinger transformed community matrix.

| Variable | $r^2$ | $\Pr(>r)$ |
| --- | --- | --- |
| Slope | 0.188 | 0.136 |
| Lithology | 0.385 | 0.001 |
| Geomorphology | 0.265 | 0.009 |
| Forest loss | 0.071 | 0.264 |

## 4 Discussion

We hypothesized that there is spatial variation in the focal Caribbean dry forest typified by clusters of plants associated with lithology, geomorphology and slope. We confirm that there is substantial spatial heterogeneity in the plant community composition of semi-deciduous forests in the Dominican Republic. Specifically, we found support for the hypothesis that plant species significantly associate with sites characterized by two qualitative factors, lithology and geomorphology. Accordingly, our findings provide new evidence of the links between geodiversity and biodiversity in tropical dry forests, and support the hypothesis association between forest communities and site groups characterized by lithological attributes found in previous research [37]. Below we discuss a selection of the detected species-habitats associations.

Several studies support most of the species-habitats associations we detected in our study. For example, *Phyllostylon rhamnoides* (Ulmaceae) and *Savia sessiliflora* (Phyllanthaceae) often occurs on calcereous soils within dry tropical forests [21, 47, 55]. Similarly, *Acacia skleroxyla* (Leguminosae) is considered a calciphilous species (i.e., adapted to life in calcium-rich and clayish soils typically developed on marlstone), and *Randia aculeata* (Rubiaceae) is commonly reported in fertile soils developed on limestone substrate [11, 12, 21, 29]. Moreover, *Hura crepitans* (Euphorbiaceae), *Picramnia pentandra* (Picramniaceae) and *Trichilia pallida* (Meliaceae), are categorized as hygrophilous species, or occuring in riparian forests on alluvial soil [6, 21, 48].

Although we found no significant association between plants and forest loss following the passive *post hoc* explanation fitting method, we do recognize that both *IndVal* and *r*_*pb*_ calculations suggest a strong association with forest loss for certain species. For example, *Coccothrinax argentea* (Arecaceae), an endemic palm, is strongly associated with sites which experienced recent tree cover loss (group D), which is in line with this species’ needs for direct sunlight and well-drained soils [27]. Similarly, the presence of a species such as *Leucaena leucocephala* (Leguminosae) in this group–though present at all groups of sites it is particularly associated with group D–can easily be explained, as this agrees with the notion that this is an invasive species that quickly becomes abundant under open canopies (e.g., elsewhere it is known to invade pastures) [16, 18, 34, 59]. More surprisingly, we also found some species that were not previously considered associated with forest disturbances or conditions of direct sunlight occurring in group D, such as *Ateleia gummifera* and *Pictetia sulcata* (both Leguminosae), which suggests that these species may be related to disturbed forests or calcium-rich soils [21].

Finally, as concluding remarks, we highlight the significance of geodiversity attributes to establish a set of habitat types. The *Random Forest* algorithm, in unsupervised learning mode, is a suitable method to classify sites using variables of mixed types. We indicate that our association analyses based on *IndVal* and *r*_*pb*_ indices, and our ordination results using redundancy analysis, are suitable methods for filling the gaps of information on spatial variability of species richness and composition in Caribbean dry forests.

We propose that our analyses hold potential for the development of site-specific management in these forests, and may support their conservation by restoration practices and protection of threatened semi-deciduous forests. Furthermore, we suggest that our approach can be replicated for other forests and countries in the region, and that our methods can be scaled up to inform regional conservation planning. Therefore, our findings could be applied in regional and local conservation planning efforts, specifically in restoration practices and in the selection of plant communities to maximize representativeness in protected areas [19, 52].

## Acknowledgement(s)

We thank Iris Santos, Gerónimo Laurencio, Edwin Medina, María de Aza and Wendy Rojas who aided in the field data collection. Design of the work was discussed with Rafael Cámara Artigas, from University of Seville, who also helped in the acquisition of field data. Species identification was done by Teodoro Clase, with the support of the National Botanical Garden of Santo Domingo. Geological maps were kindly facilitated by National Geological Survey of the Dominican Republic. Aerial photographs were donated by National Institute for Hydraulic Resources.

